# Speed-driven transitions between discrete and rhythmic dynamics in walking revealed by kinematic smoothness and muscle synergies

**DOI:** 10.64898/2026.04.09.717373

**Authors:** Giulia Panconi, Diego Minciacchi, Riccardo Bravi, Nadia Dominici

## Abstract

Humans control movement through motor primitives that generate discrete and rhythmic actions. We investigated whether and how speed may drive a transition between discrete and rhythmic organization in walking, and whether muscle synergy changes are associated with kinematic shifts. Eighteen healthy adults walked on a treadmill during incremental and decremental trials (0.5-5 km/h in 0.5 km/h steps). Kinematics and bilateral lower-limb EMG were recorded. Smoothness was quantified using log dimensionless jerk (LDJ) and spectral arc length (SPARC). Both metrics indicated lower smoothness at low speeds and progressively stabilized as speed increased, with a transition region around 3-3.5 km/h showing inter-individual variability. In parallel, EMG synergies showed speed-dependent increases in dimensionality (2→3→4), consistent with module merging at slower speeds. Overall, these findings reveal coordinated kinematic and neuromuscular shifts with speed, indicating a transition from a discrete-dominated regime at low speeds toward a more stable rhythmic pattern at higher speeds.

## INTRODUCTION

An increasing consensus suggests that human motor control is organized through a combination of fundamental building blocks that consist of dynamic primitives (Flash and Hochner, 2005; Sing et al., 2009; Degallier and Ijspeert, 2010; Dominici et al., 2011; Hogan and Sternad, 2013). Within this framework, primitives provide a model of how the central nervous system generates and controls motor behavior and, at the end-effector level, give rise to discrete and rhythmic movements (Hogan and Sternad, 2012, 2013). Discrete movements are modeled as point-attractor dynamics (or submovements) and are characterized by transitions between distinct stable postural endpoints, whereas rhythmic movements are modeled as limit-cycle dynamics that produce periodic oscillations with stable phase relationships (Hogan and Sternad, 2007, 2012, 2013; Goto et al., 2014).

Multiple lines of evidence support the theory that discrete and rhythmic movements recruit partly distinct neural networks. Neuroimaging studies have shown that discrete actions engage broader bilateral and non-primary motor areas relative to rhythmic ones, implicating partly separable cortical and cerebellar networks (Schaal et al., 2004; Spencer et al., 2007). At the behavioral level, motor learning studies have shown asymmetric transfer and reduced interference between discrete and rhythmic contexts, consistent with partly separable motor mechanisms (Ikegami et al., 2010; Howard et al., 2011). This distinction is also reflected in clinical populations, where rhythmic arm movements appear less impaired than discrete ones following stroke (Leconte et al., 2016).

Importantly, this evidence does not imply a rigid dichotomy between movement classes. Research on upper-limb control has shown that producing very slow rhythmic movements is challenging and may fragment into a sequence of discrete submovements, suggesting that the motor system can shift flexibly between these two modes as task constraints change (Sternad et al., 2013; Park et al., 2017). Although most empirical evidence supporting the discrete–rhythmic framework derives from upper-limb movements, dynamic primitives are assumed to reflect general organizational principles of the central nervous system, and therefore similar transitions should in principle be observable across different effector systems, including locomotion.

Locomotion provides a particularly relevant context to test this idea. Walking is typically considered a strongly rhythmic behavior, yet at very slow speeds coordination may become less stable and exhibit features resembling sequences of discrete submovements. This makes locomotion a particularly interesting model to investigate how the motor system transitions between discrete-like and rhythmic organization under changing task constraints. This is particularly relevant as locomotor speed is a key variable in both daily life and clinical contexts. Slow walking is common in aging, neurological conditions, and early rehabilitation, precisely conditions in which rhythmic locomotor organization may be compromised.

Walking speed has been proposed as a control parameter triggering changes in coordination patterns consistent with phase-transition-like behavior (Diedrich and Warren, 1995, 1998; Raffalt et al., 2020). Building on this, Moura Coelho et al. (2022) showed that kinematic smoothness measures can serve as a proxy for distinguishing discrete-like from rhythmic-like walking patterns, with very slow walking exhibiting stronger discrete-like features and faster walking reflecting a more stable rhythmic organization. These findings suggest that locomotion may not be uniformly rhythmic across speeds, but rather operate within a range of regimes whose expression depends on task constraints. Together, these findings indicate that speed-dependent regime shifts may occur in locomotion, although the mechanisms driving these shifts remains poorly understood.

Despite the increasing evidence, key questions remain regarding how discrete-rhythmic regime shifts evolve across a range of walking speeds and whether consistent patterns emerge across individuals. Furthermore, the existing evidence within the discrete-rhythmic primitives literature remains predominantly kinematic, leaving open whether kinematic regime shifts are accompanied by systematic reorganization of neuromuscular activity.

Muscle synergy analysis provides a strong framework to characterize the modular organization of locomotor muscle activity, where groups of muscles are co-activated in coordinated patterns that can be flexibly recruited across task conditions (Cheung et al., 2005; Ivanenko et al., 2005). Importantly, EMG-based studies have shown that distinct spinal locomotor networks can be differentially recruited across locomotor modes and that muscle activation patterns may reorganize with speed even within the same locomotor mode (Yokoyama et al., 2016), raising the possibility that transitions observed at the kinematic level may reflect underlying changes in the organization of neuromuscular control.

Accordingly, this study aims to characterize the speed-dependent reorganization between discrete-like and rhythmic-like locomotor patterns, focusing on how locomotor control reorganizes across walking speeds. Specifically, we examine whether this transition occurs gradually or exhibits a more pronounced shift within a particular speed range, quantify inter-individual consistency, and determine whether kinematic regime shifts are accompanied by systematic EMG reorganization assessed through changes in muscle synergies, thereby linking changes in movement dynamics to their underlying control mechanisms.

## METHODS

### Participants

18 healthy adults were recruited as participants for this study (12 female and 6 male, age 21.0 ± 2.5 years [mean ± SD], height 172.7 ± 8.8 cm, weight 65.7 ± 11.7 kg; see Table S1 for individual participant characteristics). A priori power analysis for a repeated-measures ANOVA (within-subjects design; power = 0.8, α = 0.05, effect size f = 0.25, one group) indicated that a minimum of 14 participants were required to detect statistically meaningful effects. We intentionally set the target recruitment above this threshold (n = 18) to account for inter-individual variability, potential attrition or unusable data due to technical issues. Exclusion criteria included the presence of neurological or musculoskeletal disorders, as well as recent, ongoing, or permanent injuries that may influence locomotor function and alter typical gait patterns.

The study was conducted in accordance with the guidelines of the Declaration of Helsinki. All procedures were approved by the Scientific and Ethical Review Board of the Faculty of Human Movement Sciences, Vrije Universiteit Amsterdam (VCWE-2025-119). Before the experiment, all participants read and signed a written informed consent form.

### Experimental Design

Prior to the main experimental conditions, participants completed a familiarization phase on the treadmill to acclimate to walking at varying speeds. The main experimental protocol consisted of participants walking on an instrumented treadmill (N-Mill, 60 × 150 cm, Motek Medical B.V., Amsterdam, the Netherlands) under two conditions: incremental and decremental speed. Each participant performed one trial under each condition. In the incremental speed condition, participants started walking at 0.5 km/h, and the speed was increased in steps of 0.5 km/h until reaching 5 km/h. In the decremental speed condition, participants started walking at 5 km/h, and the speed was decreased in steps of 0.5 km/h until reaching 0.5 km/h. At each speed, participants completed 15 consecutive strides before the speed was changed. Short rest breaks were allowed when needed to minimize fatigue. The order of the incremental and decremental trials was randomized across participants.

### Data Acquisition

Kinematic data were recorded using a Vicon motion capture system (Vicon Vero v2.2, Oxford, UK) consisting of 10 infrared cameras placed around the treadmill and sampling at 100 Hz (Figure 1). A total of 19 infrared reflective markers of 14 mm diameter were positioned bilaterally on anatomical landmarks: heels, fifth metatarsophalangeal joints, lateral malleoli, lateral femur epicondyles, greater trochanters, iliac spinal crests, glenohumeral joints, humeral epicondyles, and ulnar styloids. An additional Vicon Vue video camera was positioned in the participants’ right sagittal plane at 100 Hz.

**Figure 1.**
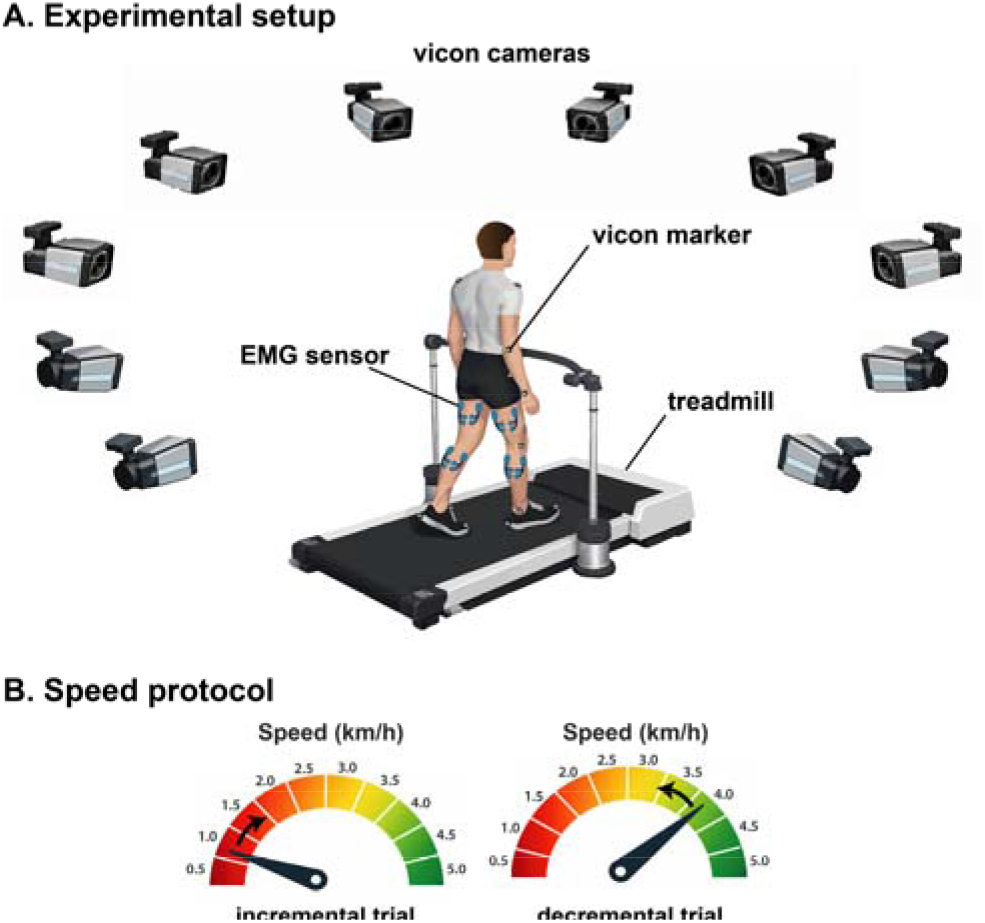
**A. Experimental setup**. Participants walked on a treadmill while kinematics were recorded with a 10-camera Vicon motion capture system. Reflective markers were placed bilaterally on anatomical landmarks (upper limb, pelvis, and lower limb), and EMG activity was recorded bilaterally: from lower-limb muscles. **B. Speed protocol**. Participants performed an incremental trial with speed increased from 0.5 to 5.0 km/h in 0.5 km/h steps, and a decremental trial with speed decreased from 5.0 to 0.5 km/h in 0.5 km/h steps.

Muscle activity was recorded using a Wave Plus wireless EMG system with mini probes (Cometa, Milan, Italy) with Ag/AgCl electrodes (Ambu Blue Sensor). EMG signals were collected from 15 muscles bilaterally: tibialis anterior (TA), gastrocnemius lateralis (GL), gastrocnemius medialis (GM), soleus (SOL), peroneus longus (PL), rectus femoris (RF), vastus lateralis (VL), vastus medialis (VM), sartorius (SART), biceps femoris (BF), semitendinosus (SEM), tensor fasciae latae (TFL), gluteus maximus (GLM), gluteus medius (GLMED) and erector spinae at the level of L2 (ES).

EMG signals were amplified with a gain of 1,000, and online band-pass filtered between 10 and 500 Hz before sampling at 2000 Hz.

In preparation, the participants’ skin was cleaned with alcohol at the locations where the electrodes were attached. Electrodes were attached to the skin over the muscle belly according to standard recommendations for minimizing crosstalk between adjacent muscles (Hermens et al., 2000). Once all markers and electrodes were applied directly to the skin, participants’ clothing was adjusted to ensure that they did not contact or interfere with either the markers or the electrodes. EMG and kinematics were simultaneously acquired and synchronized.

### Kinematic preprocessing and event detection

Data were processed using MATLAB custom script (2024b The MathWorks, Natick, MA, USA). Marker trajectories were extracted and low-pass filtered at 10 Hz (bidirectional 2nd-order Butterworth filter). Heel-strike (HS) and toe-off (TO) events were detected for both limbs using peak-based distance metrics (Zeni et al., 2008) and subsequently verified using an interactive GUI that allowed manual inspection and correction of gait events to ensure accuracy. HS and TO events were used to compute spatiotemporal gait parameters, including step time and stride time (from successive HS), cadence (steps/min), step and stride length, and stance, swing, and double-support durations and their percentages of the gait cycle for each limb. Treadmill speed was estimated from the anterior-posterior heel displacement between consecutive gait events and was used to segment each trial into constant-speed blocks.

### Smoothness analysis

To capture inter-limb gait dynamics, we analyzed a relative ankle kinematic variable defined as the antero-posterior displacement between the right and left ankles (lateral malleoli marker) (Plotnik et al., 2007; Moura Coelho et al., 2022). The right-left ankle displacement signal was demeaned and segmented between successive zero-crossings, with each segment defined by two zero-crossings sharing the same slope sign. Movement smoothness was then quantified for each segment using two complementary smoothness metrics: Log Dimensionless Jerk (LDJ) and Spectral Arc Length (SPARC) (Balasubramanian et al., 2015, 2012; Melendez-Calderon et al., 2021). Lower smoothness values were taken to reflect a less continuous and more segmented movement pattern, whereas higher smoothness values were taken to indicate more continuous rhythmic dynamics.

Logarithmic Dimensionless Jerk (LDJ) is a time-domain metric that normalizes integrated squared jerk for movement duration and amplitude, making it largely invariant to scaling and enabling comparisons across conditions (Hogan and Sternad, 2009; Balasubramanian et al., 2012, 2015). LDJ was computed as:

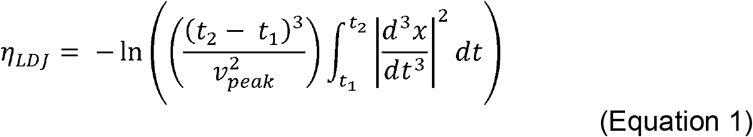

LDJ values are negative, with values closer to zero (i.e., less negative) indicating smoother movements (Hogan and Sternad, 2009; Balasubramanian et al., 2012, 2012; Gulde and Hermsdörfer, 2018).

Spectral Arc Length (SPARC) is a frequency-domain metric that quantifies smoothness from the spectral complexity of the speed profile, and has been shown to be robust to measurement noise and less sensitive to scaling effects than jerk-based measures (Balasubramanian et al., 2012, 2015; Melendez-Calderon et al., 2021). SPARC was computed as:

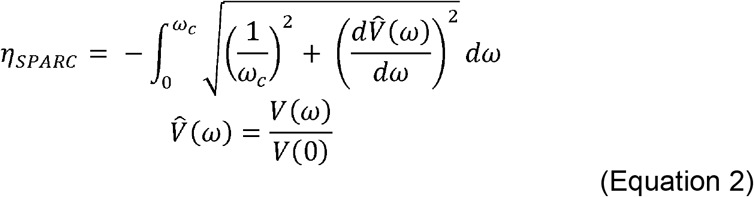

where ω_c is the upper frequency limit of integration, V(ω) is the magnitude spectrum of the speed profile, and V□(ω) is the spectrum normalized by its value at zero frequency. As with LDJ, SPARC values are negative, with values closer to zero indicating smoother movements (Balasubramanian et al., 2012, 2015). The two metrics were interpreted jointly to offset the respective limitations of time- and frequency-domain approaches.

### EMG preprocessing

EMG data were visually inspected to identify artifacts, and corrupted data segments were excluded from further analysis. On average, 3.7 ± 2.0 EMG channels were excluded per participant. The raw EMG was high-pass filtered at 30 Hz (bidirectional 4th-order Butterworth filter) to suppress low-frequency movement artifacts. Power-line noise was removed using a series of notch filters at 50, 100, and 150 Hz (bidirectional 4th-order Butterworth filter). In addition, to remove artifacts generated by the wireless accelerometer embedded in the EMG sensors a narrow band-pass filter was applied around 142 ± 2 Hz. The cleaned signal was then rectified and low-pass filtered at 10 Hz (bidirectional 4th-order Butterworth) to extract the linear envelope.

Right heel-strike events were used to segment the continuous EMG signal into individual gait cycles. Each gait cycle was time-normalized to 200 samples and amplitude-normalized to the peak EMG amplitude across the full trial.

For each stride, treadmill belt speed was estimated as the anterior-posterior velocity of the heel marker during the stance phase, reflecting the period of direct foot-belt contact. Speed-transition events identified from the kinematic segmentation were used to define the boundaries of speed-based EMG segments.

### EMG synergy extraction

Muscle synergies were extracted from the EMG data using non-negative matrix factorization (NNMF), a linear decomposition technique that approximates the EMG data matrix as the product of two non-negative matrices representing time-invariant muscle weightings and time-varying activation profiles (Lee and Seung, 1999; Cheung et al., 2005). Specifically, the EMG matrix 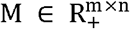 (with muscles and n time samples) was factorized as:

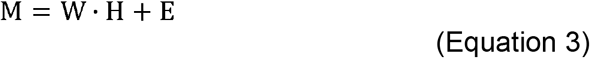

where 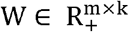 contains the synergy weight vectors (spatial modules or muscles weightings), 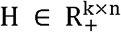 contains the corresponding activation coefficients (temporal profiles or activation patterns), K is the number of synergies, and E is the residual error matrix. Equivalently, the EMG data can be expressed as a non-negative linear combination of K synergies:

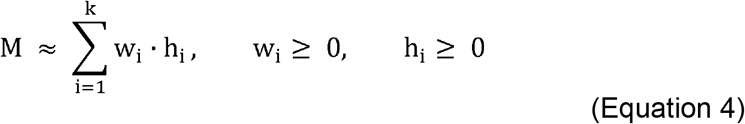

Synergy analysis was performed separately for incremental and decremental conditions using two parallel pipelines: a subject-level NNMF and a group-level NNMF. At the subject level, EMG data were organized into speed-defined segments and time-normalized over the gait cycle; within each speed-segment, the cycle-averaged EMG matrix (mean across gait cycles) was factorized into W and H. The number of synergies k was varied from 1 to 10; for each speed segment, NNMF was run 100 times with random initializations, with a maximum of 1,000 iterations per run, and the solution yielding the lowest reconstruction error was retained. Factorization was performed using the multiplicative update algorithm. Reconstruction quality was quantified as variance accounted for (VAF), computed as VAF = 1− SSE / SST, where SSE is the sum of squared reconstruction errors and SST is the total sum of squares of the original EMG data. The final number of synergies was selected using the 90% VAF criterion (smallest k such that VAF ≥ 0.90) (Torres-Oviedo et al., 2006; Hashiguchi et al., 2016; Yokoyama et al., 2016).

In parallel, at the group level, a segment-wise group EMG matrix was constructed by averaging each subject’s cycle-averaged EMG profile within each speed segment, resulting in a single 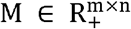 matrix for each segment and condition. The same NNMF procedure (k = 1–10; 100 runs per k; 1000 iterations, 90% VAF criterion) was applied to obtain group-representative synergy structures. Because NNMF components are not inherently ordered, synergies were first ordered by the timing of their peak activation in H in a reference (“leader”) segment (5 km/h) and then matched across segments using a similarity criterion to ensure consistent labeling across speeds.

### Merging analysis

To examine whether changes in synergy dimensionality across walking speeds could be explained by merging of the modules, a reconstruction-based procedure was implemented following Cheung et al., (2020). In this approach, synergy merging refers to the process by which distinct synergies active at higher speeds are subsumed into fewer synergies at lower speeds; under this model, each slower-speed synergy can be approximated as a non-negative linear combination of faster-speed synergies. Evidence for merging was therefore inferred when synergies from the faster segment successfully reconstructed those from the slower speed-segment, consistent with a reduction in synergy number driven by the combining of modules across speeds.

For each trial type, the group-averaged synergy weight matrices (W) estimated within each speed segment were analyzed. Merging was evaluated across adjacent speed segments in the direction faster → slower, such that synergies from the faster segment were used as basis vectors to reconstruct synergies from the slower segment. Prior to fitting, all synergy weight vectors were normalized to unit length (Cheung et al., 2020).

For each target synergy 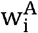 in the slower speed-segment A (segA), non-negative coefficients *m*_*ik*_ were estimated to best reconstructed the target from the faster speed-segment synergies 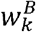 in segment B (segB) using non-negative least squares (MATLAB function lsqnonneg). The reconstruction model was:

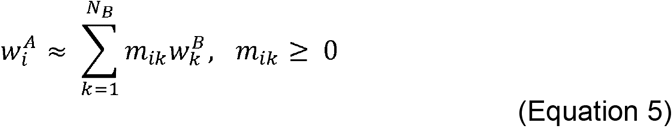

To retain only meaningful contributors, a threshold of m_ik_ ≥ 0.20 was applied to the estimated coefficients, as defined in Cheung et al., (2020).

A target synergy was classified as merged when it was reconstructed by at least two contributing synergies above this threshold and the cosine similarity between the target and its reconstruction was at least 0.80 (Cheung et al., 2020). For each transition, merging was summarized as the count and proportion of target synergies in the slower speed-segment meeting these criteria.

### Statistical analysis

In order to quantify whether smoothness parameters systematically changed across speed-defined segments within individual trials, a within-trial repeated-measures analysis was conducted separately for each subject and condition. A repeated-measures ANOVA with segment as the within-trial factor was used to test for an overall speed segment effect. Where a significant effect was found, pairwise post-hoc comparisons between segments were extracted from the fitted repeated-measures model, with p-values adjusted for multiple comparisons using the Holm adjustment, which was preferred for its balance between control of familywise error rate and statistical power.

In the present study, the transition was interpreted as the speed range over which smoothness measures showed their clearest reorganization across speed segments, rather than as a single threshold.

To examine whether the number of extracted muscle synergies systematically changed across speed-defined segments, a within-subject repeated-measures analysis was conducted separately for the incremental and decremental conditions. For each subject, the number of synergies identified per speed segment served as the unit of observation. A one-factor repeated-measures ANOVA with speed as the within-subject factor was used to test for an overall speed effect. Pairwise post-hoc comparisons between speed levels were obtained from the fitted model, with p-values adjusted for multiple comparisons using the Holm adjustment.

To visualize the consistency of speed-related effects across participants, the results of the pairwise post-hoc comparisons were summarized as the percentage of participants showing a significant difference (Holm-adjusted p < 0.05) for each speed pair. These percentages were visualized as heatmaps to illustrate the distribution of significant contrasts across walking speeds.

## Results

### Speed-dependent changes in kinematic smoothness

Both smoothness metrics showed a clear dependence on walking speed, with greater variability at low walking speeds and progressively more stable values at the highest speeds (Figures 2 and 3).

**Figure 2.**
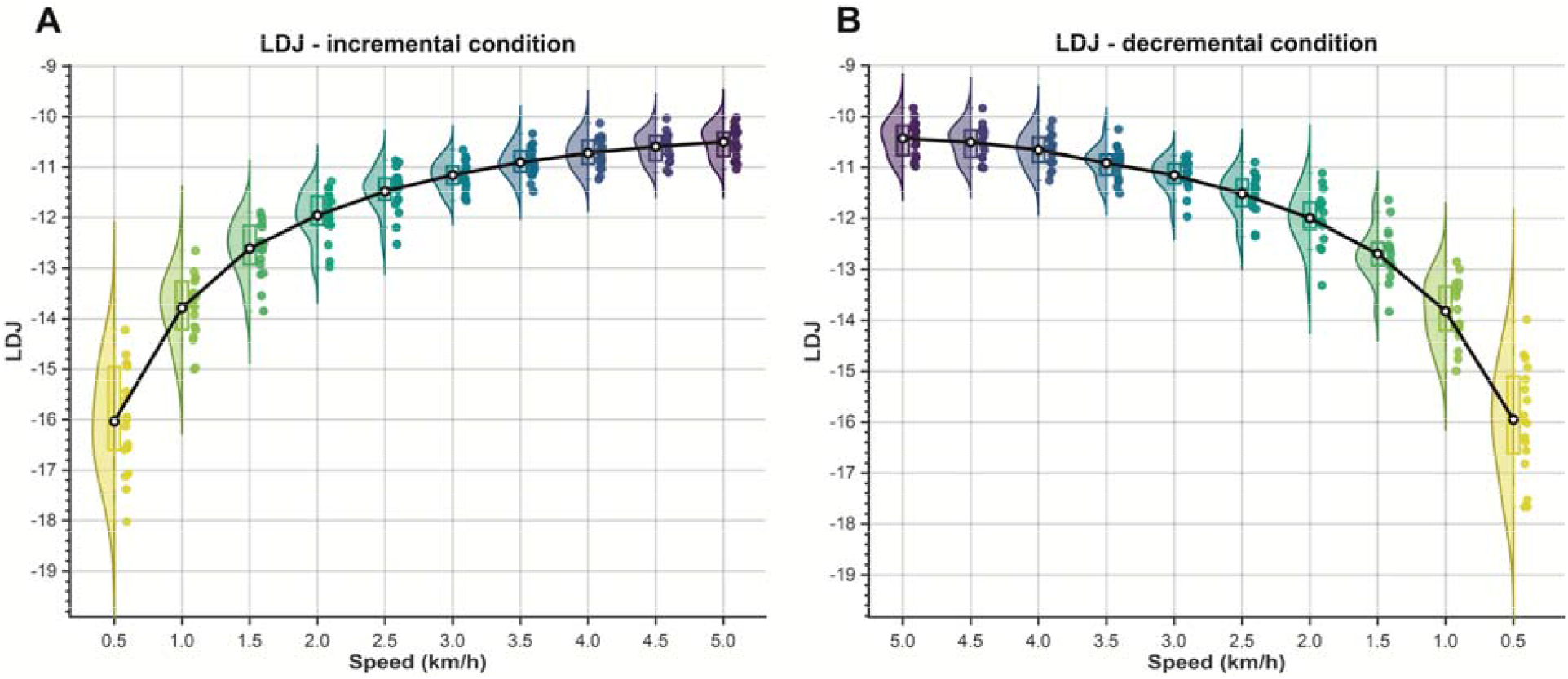
Speed-dependent changes in LDJ during incremental (A) and decremental (B) conditions. The distribution of LDJ values computed for each speed segment for the incremental condition (A) and the decremental condition (B). Colored points represent each participant’s mean at that speed segment, and the dots connected by a black line indicate the group mean across participants. LDJ: logarithmic dimensionless jerk.

**Figure 3.**
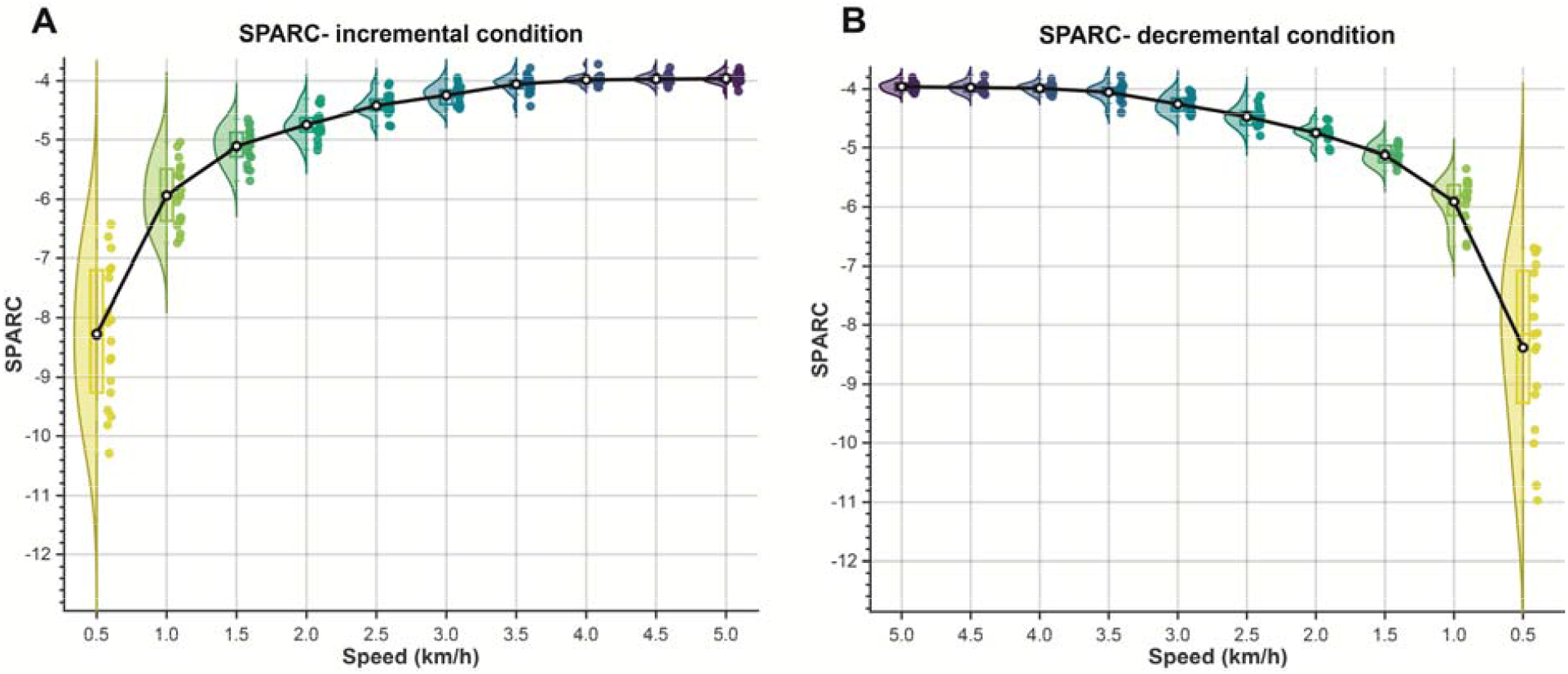
Speed-dependent changes in SPARC during incremental (A) and decremental (B) conditions. The distribution of SPARC values computed for each speed segment for the incremental condition (A) and the decremental condition (B). Colored points represent each participant’s mean at that speed segment, and the black dot connected by a black line indicates the group mean across participants. SPARC: spectral arc length.

LDJ values increased (i.e., became less negative) with speed in both conditions, with mean subjects values increasing from −16.03 ± 1.14 and −15.95 ± 1.20 at 0.5 km/h (incremental and decremental, respectively) to −10.50 ± 0.34 and −10.43 ± 0.36 at 5 km/h. (Figure 2).

A similar pattern was observed for SPARC, with values becoming progressively less negative as speed increased, from −8.28 ± 1.55 and −8.38 ± 1.71 at 0.5 km/h (incremental and decremental, respectively) to −3.96 ± 0.09 and −3.96 ± 0.08 at 5 km/h.

Within-trial repeated-measures ANOVA revealed a significant effect of speed on LDJ across subjects (p<0.001) (Figure 4). Pairwise post-hoc comparisons revealed highly consistent LDJ differences at low-to-moderate speeds: from 0.5 to 2.5 km/h, comparisons involving lower-speed segments showed high consistency across participants (typically ≥89–100%). This consistency dropped to a moderate range around 3–3.5 km/h (approximately 50–80%, depending on the specific contrast), and became low at the highest speeds (4–4.5 km/h), where only a small fraction of subjects showed significant differences (0–11%) between high-speed segments (Figure 4).

**Figure 4.**
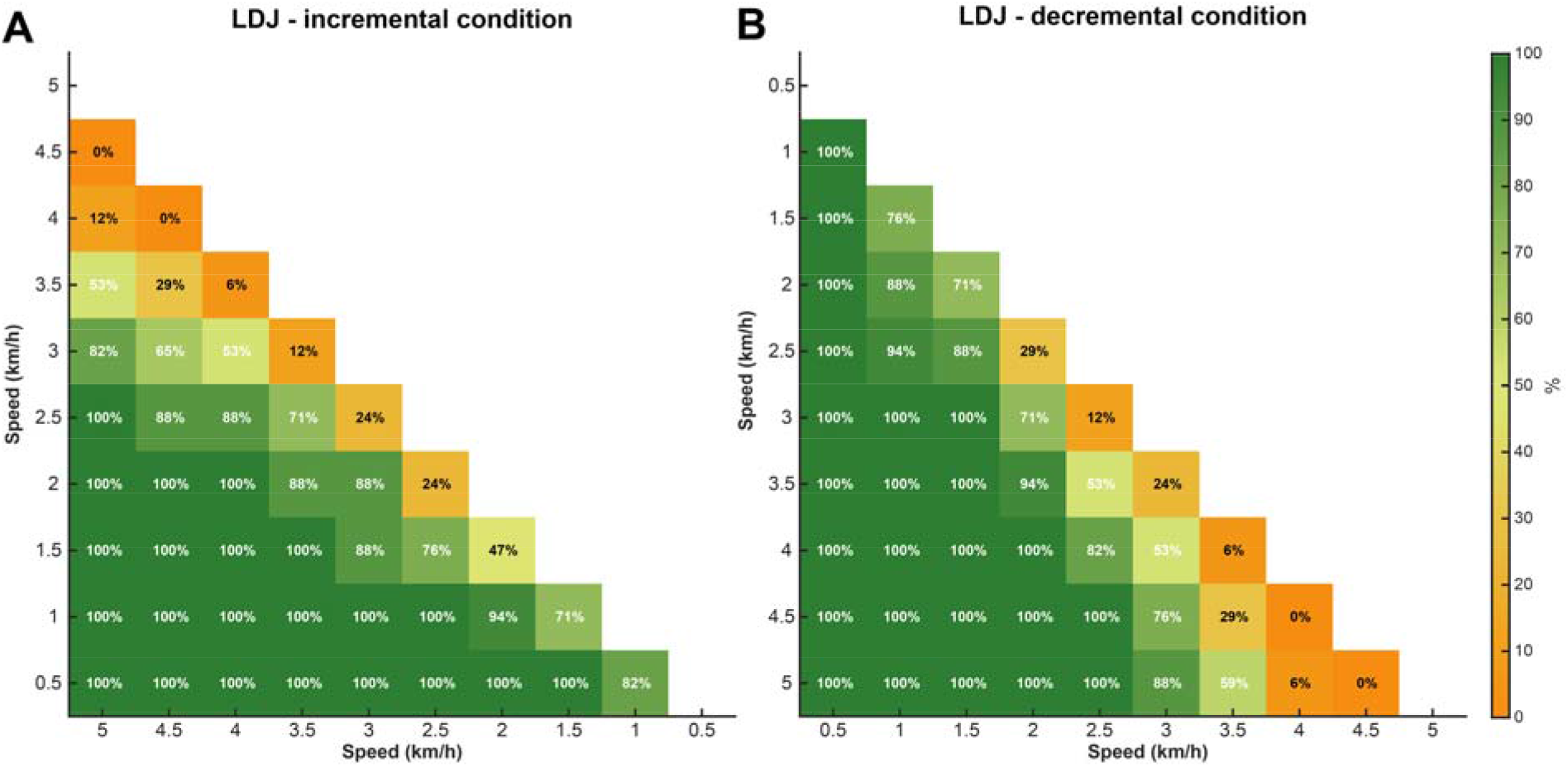
Heatmaps showing the percentage of participants with significant LDJ pairwise differences between speed segments. In incremental (A) and decremental (B) conditions, the percentage of participants showing a statistically significant difference in LDJ between each pair of walking speeds (0.5-5.0 km/h in 0.5 km/h steps). Each cell reports the % of participants with a significant pairwise contrast for the corresponding speed combination (axes), with color indicating the proportion (orange = lower, green = higher). Higher percentages indicate speed pairs that more consistently elicited LDJ changes across individuals. LDJ: log dimensionless jerk.

Within-trial repeated-measures ANOVA showed a consistent significant effect of speed on SPARC across participants (p<0.001) (Figure 5). Pairwise post-hoc comparisons revealed highly consistent SPARC differences at low-to-moderate speeds: from 0.5 to 2.5 km/h, comparisons involving lower-speed segments showed high consistency across participants (≥89%). This consistency dropped to a moderate-to-high range around 3 km/h (approximately 80%) and further decreased to low-to-moderate levels around 3.5 km/h (approximately 20–30%). At the highest speeds (4–5 km/h), agreement was very low, with no participants showing significant differences between segments (Figure 5).

**Figure 5.**
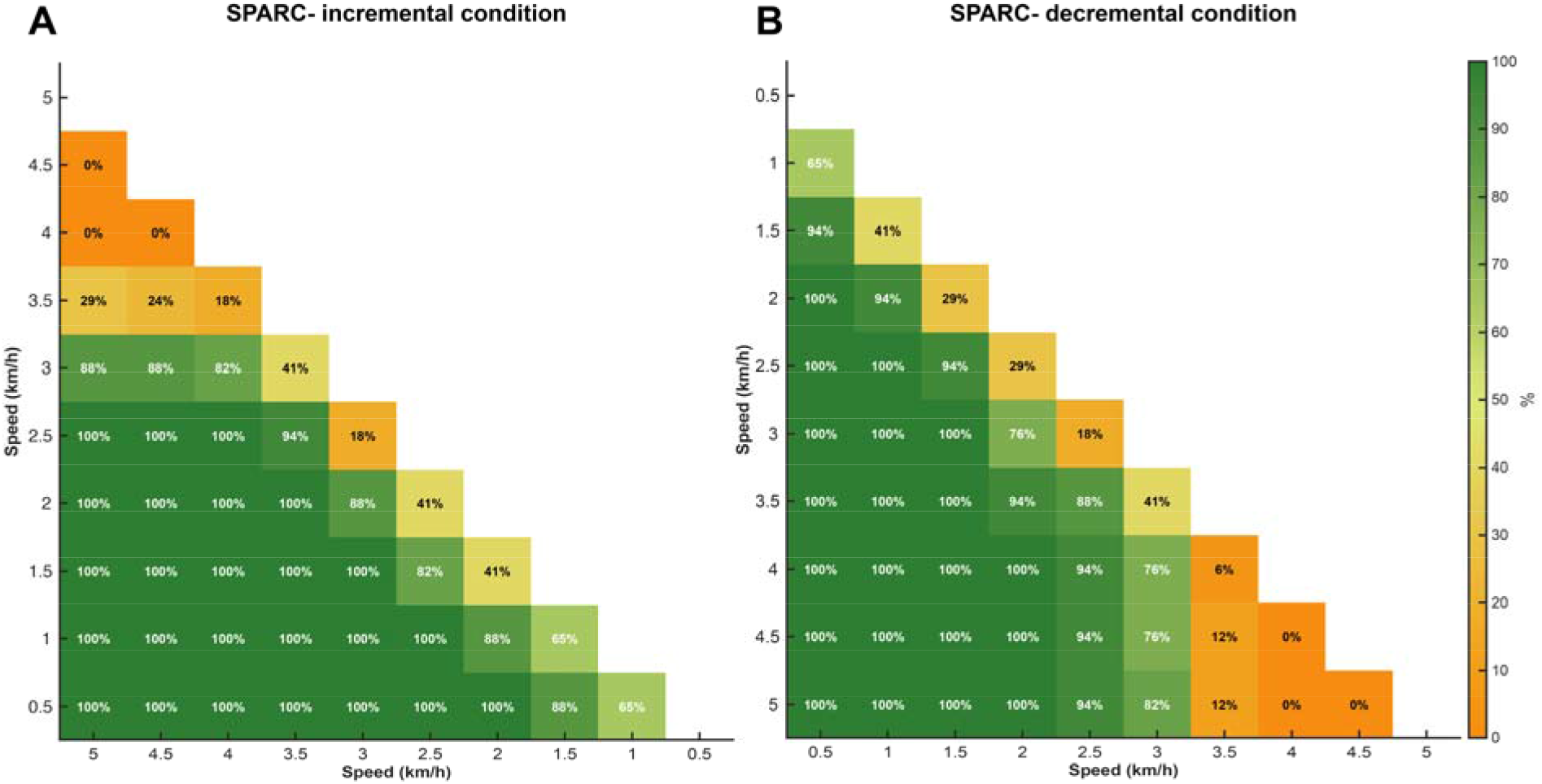
Heatmaps showing the percentage of participants with significant SPARC pairwise differences between speed segments. In incremental (A) and decremental (B) conditions, the percentage of participants showing a statistically significant difference in SPARC between each pair of walking speeds (0.5-5.0 km/h in 0.5 km/h steps). Each cell reports the % of participants with a significant pairwise contrast for the corresponding speed combination (axes), with color indicating the proportion (orange = lower, green = higher). Higher percentages indicate speed pairs that more consistently elicited SPARC changes across individuals. SPARC: spectral arc length.

### Speed-dependent modulation of muscle synergy dimensionality

Across speeds, the number of muscle synergies identified at the participant level showed a clear speed-dependent modulation in both conditions (mean ± SD across participants) (Figure 6).

**Figure 6.**
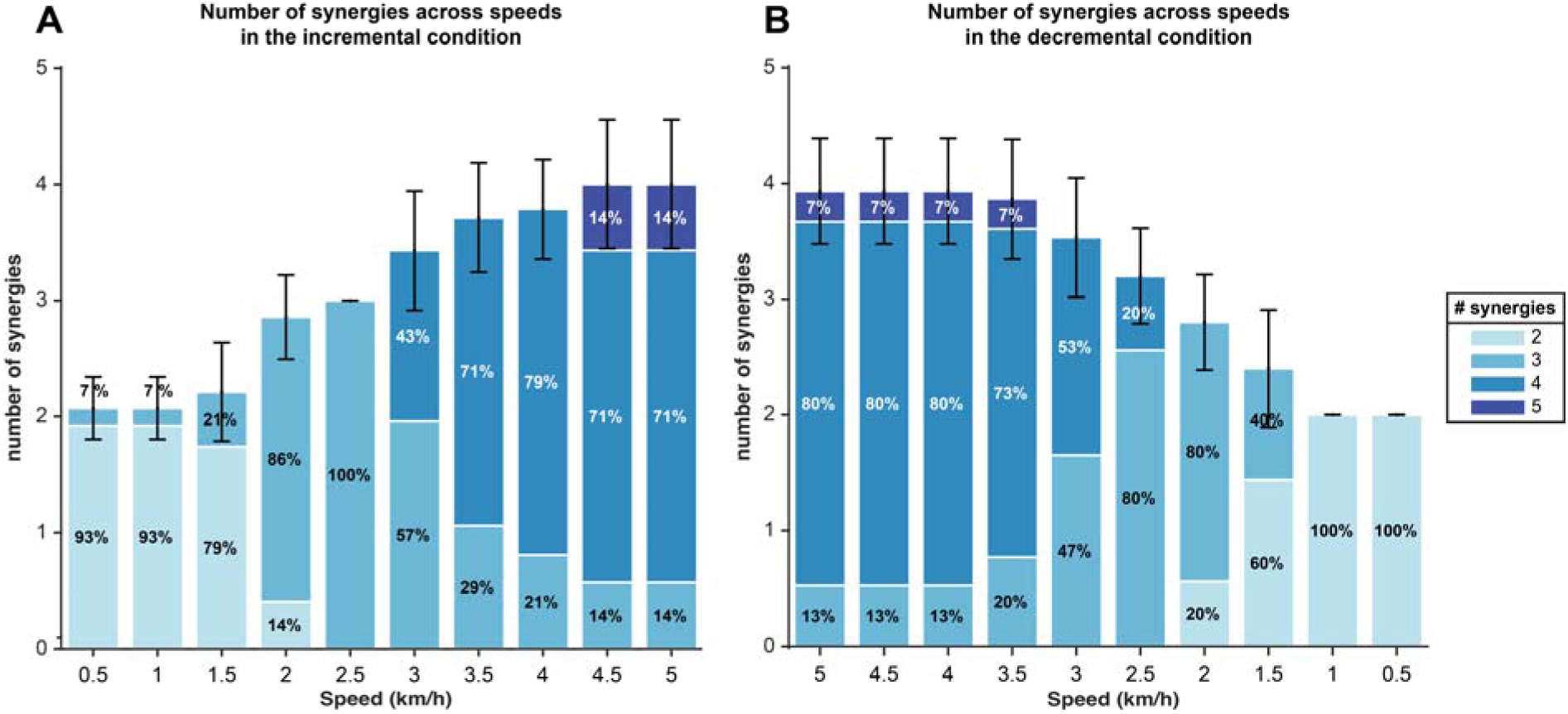
Speed-dependent distribution of the number of muscle synergies in the incremental (A) and decremental (B) conditions. Stacked bar plots show, for each speed-segment, the percentage of participants whose EMG decomposition resulted in a given number of synergies (2–5) during the incremental (A) and decremental (B) protocols. Error bars represent the between-participant standard deviation of the mean synergy number at each speed.

A one-factor repeated-measures ANOVA revealed a significant main effect of speed on the number of extracted synergies in both the incremental and decremental conditions.

In the incremental condition, the mean number of synergies increased with speed, from 2 synergies at 0.5–1.5 km/h to 3 synergies at 2–2.5 km/h, reaching 4 synergies from 3 km/h onward and stabilizing around 4 synergies at 4.5–5 km/h (Figure 6A).

In the decremental condition, the mean number of synergies was highest at fast walking (4 synergies at 5–3.5 km/h) and progressively decreased as speed was reduced, reaching ∼3 synergies at 2 km/h, and ∼2 synergies at 1–0.5 km/h (Figure 6B).

Pairwise post-hoc comparisons (Holm-adjusted) indicated a stabilization of the number of muscle synergies at higher walking speeds in both conditions. In the incremental condition, no differences were observed between 5.0 km/h and 4.5, 4.0, or 3.5 km/h (all p ≥ 0.441). In contrast, significant differences emerged when comparing the high-speed segments with 3.0 km/h (5.0 vs 3.0 km/h: p = 0.018; 4.5 vs 3.0 km/h: p = 0.018) (Figure 7A).

**Figure 7.**
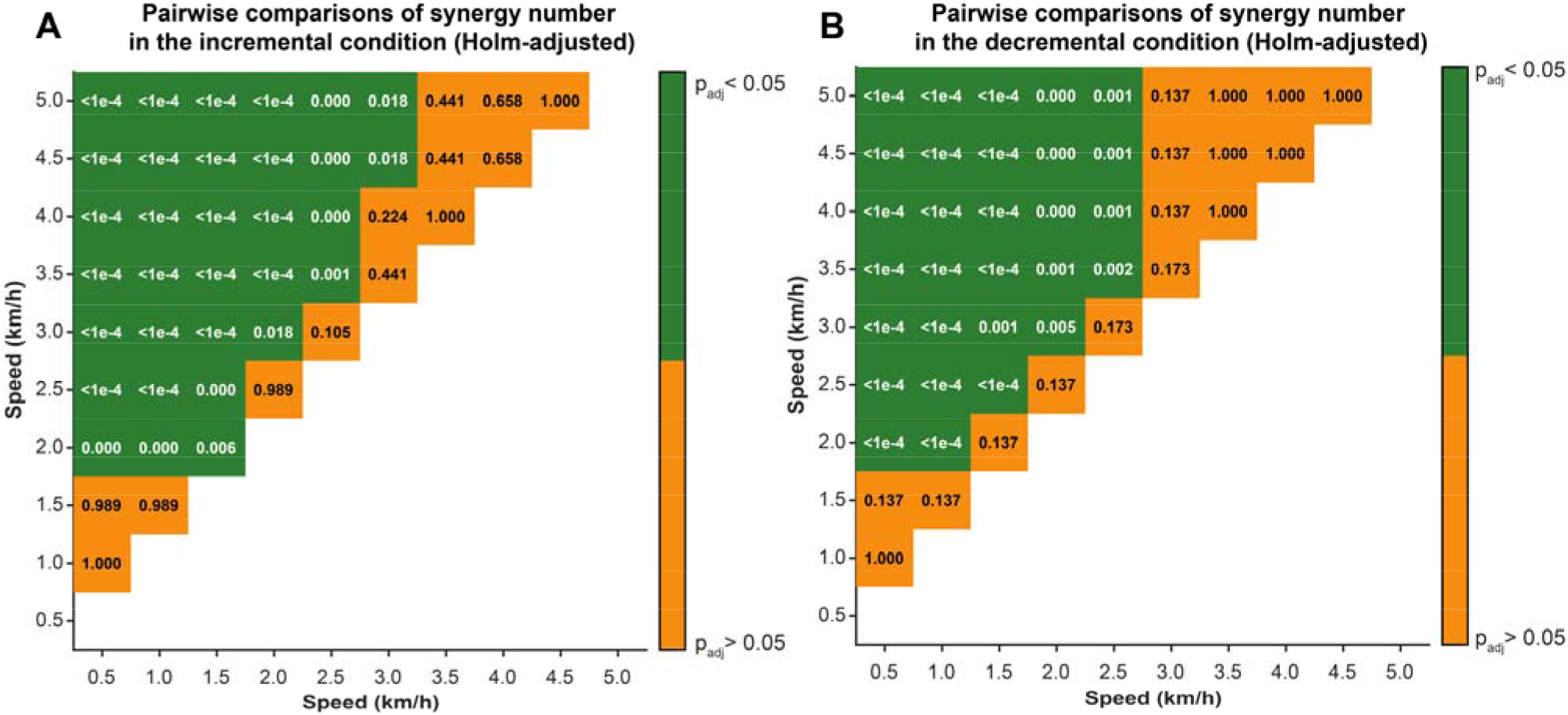
Heatmaps showing post-hoc pairwise differences in synergy number across walking speeds in incremental (A) and decremental (B) conditions. Each cell reports the Holm-adjusted p-value for the pairwise comparison between the two speed segments (km/h); green indicates significant differences (p_adj_ < 0.05), and orange indicates non-significant comparisons (p_adj_ ≥ 0.05). In both conditions, comparisons among the highest speeds show non-significant differences, whereas significant differences emerge when contrasting high-speed segments with lower-speed segments from 3 km/h in the incremental condition and 2.5 km/h in the decremental condition.

In the decremental condition, the number of synergies at 5.0 km/h did not differ from 4.5, 4.0, 3.5, or 3.0 km/h (all p ≥ 0.137). The first significant difference appeared at 2.5 km/h where the number of synergies differed from high-speed segments (5.0, 4.5, and 4.0 vs 2.5 km/h: p = 0.001) (Figure 7B).

At the group level, the EMG signals used for synergy extraction were averaged across participants within each speed-segment and condition (incremental and decremental). VAF curves were then computed from the resulting group-averaged EMG matrices for each speed-segment as a function of the number of extracted synergies in order to determine the minimum number of synergies meeting the VAF criterion (Figure 8). The curves show that higher walking speeds required a greater number of synergies to reach the VAF criterion.

**Figure 8.**
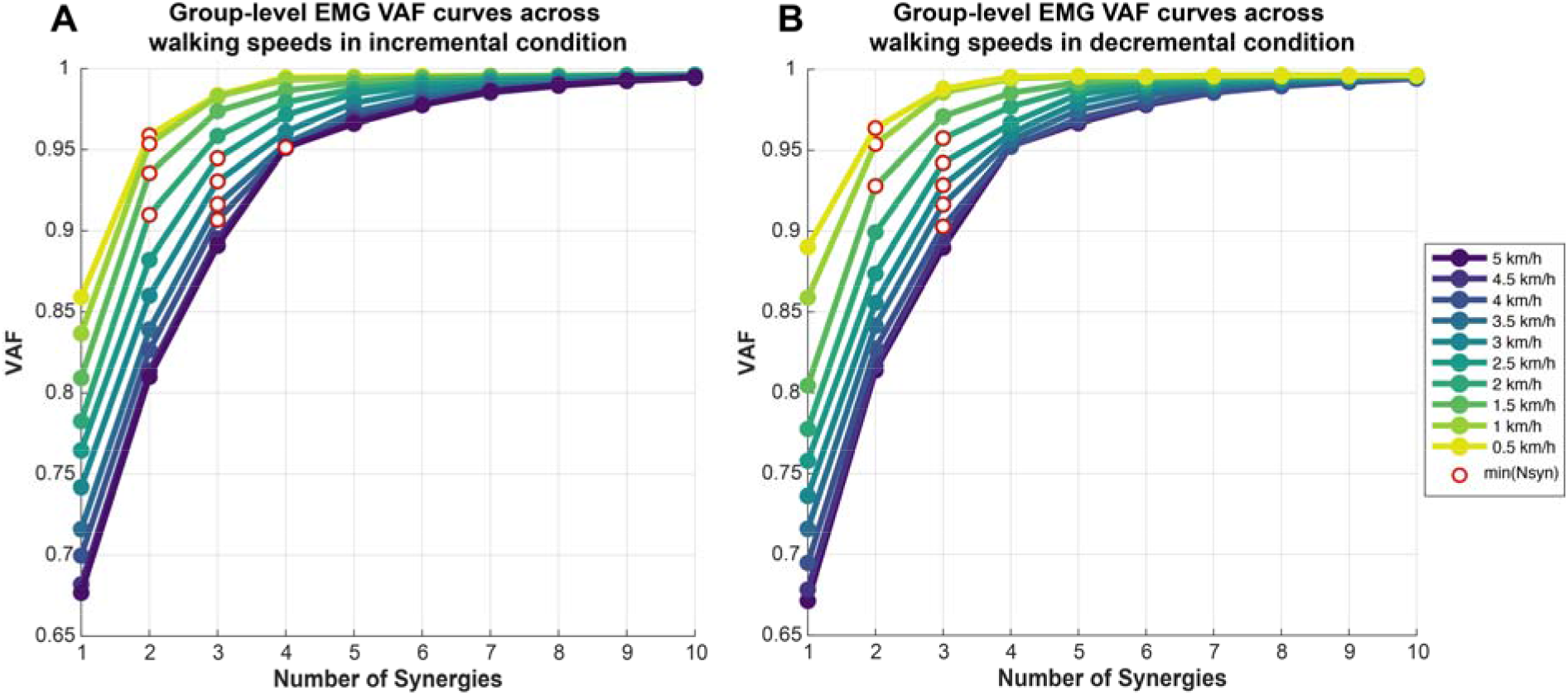
Group-level EMG VAF curves across walking speeds in the incremental (A) and decremental (B) conditions. Variance accounted for (VAF) of the reconstructed EMG signal is plotted as a function of the number of extracted muscle synergies (1–10) for each speed segment (0.5–5 km/h; speed color-coded). Open red circles indicate the selected minimal number of synergies (min(*N*syn)) at each speed according to the adopted VAF criterion (VAF≥ 0.90).

The number of synergies required to explain EMG variance showed a clear dependence on walking speed in both conditions (Figures 9 and 10).

**Figure 9.**
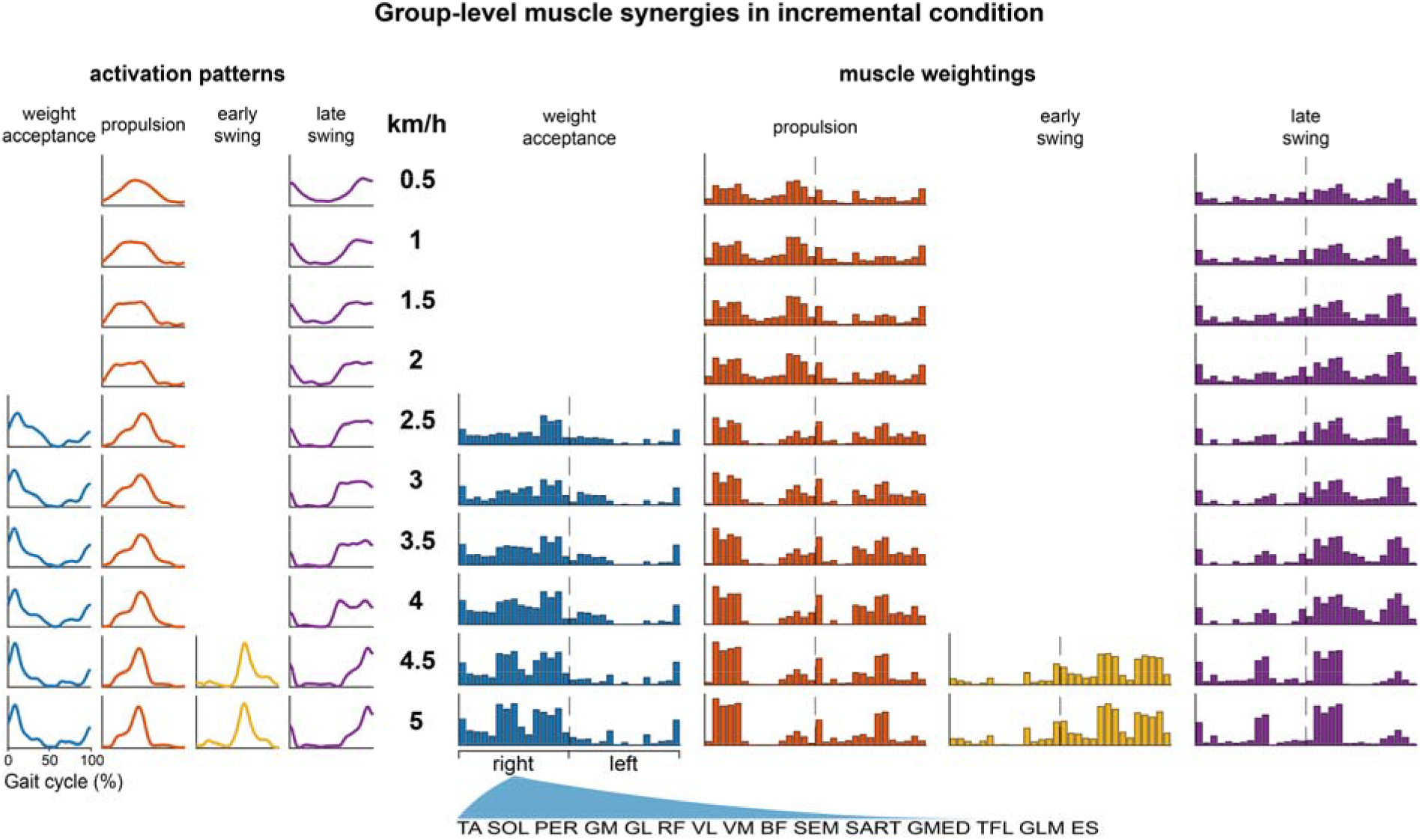
Group-level muscle synergies across speeds in the incremental condition. For each speed segment (0.5–5.0 km/h; rows), the figure shows the extracted synergy activation patterns (H; line plots on the left) and the corresponding muscle weightings (W; bar plot on the right). Activation patterns are time-normalized to the gait cycle (0-100%) and organized according to the functional phase of the gait cycle (weight acceptance, propulsion, early swing, and late swing). Bar plots show normalized muscle weights, with dashed vertical lines separating right- and left-side muscle groups. TA: tibialis anterior; SOL: soleus; PER: peroneus longus; GM: gastrocnemius medialis; GL: gastrocnemius lateralis; RF: rectus femoris; VL: vastus lateralis; VM: vastus medialis; BF: biceps femoris; SEM: semitendinosus; SART: sartorius; GMED: gluteus medius; TFL: tensor fasciae latae; GLM: gluteus maximus; ES: erector spinae.

**Figure 10.**
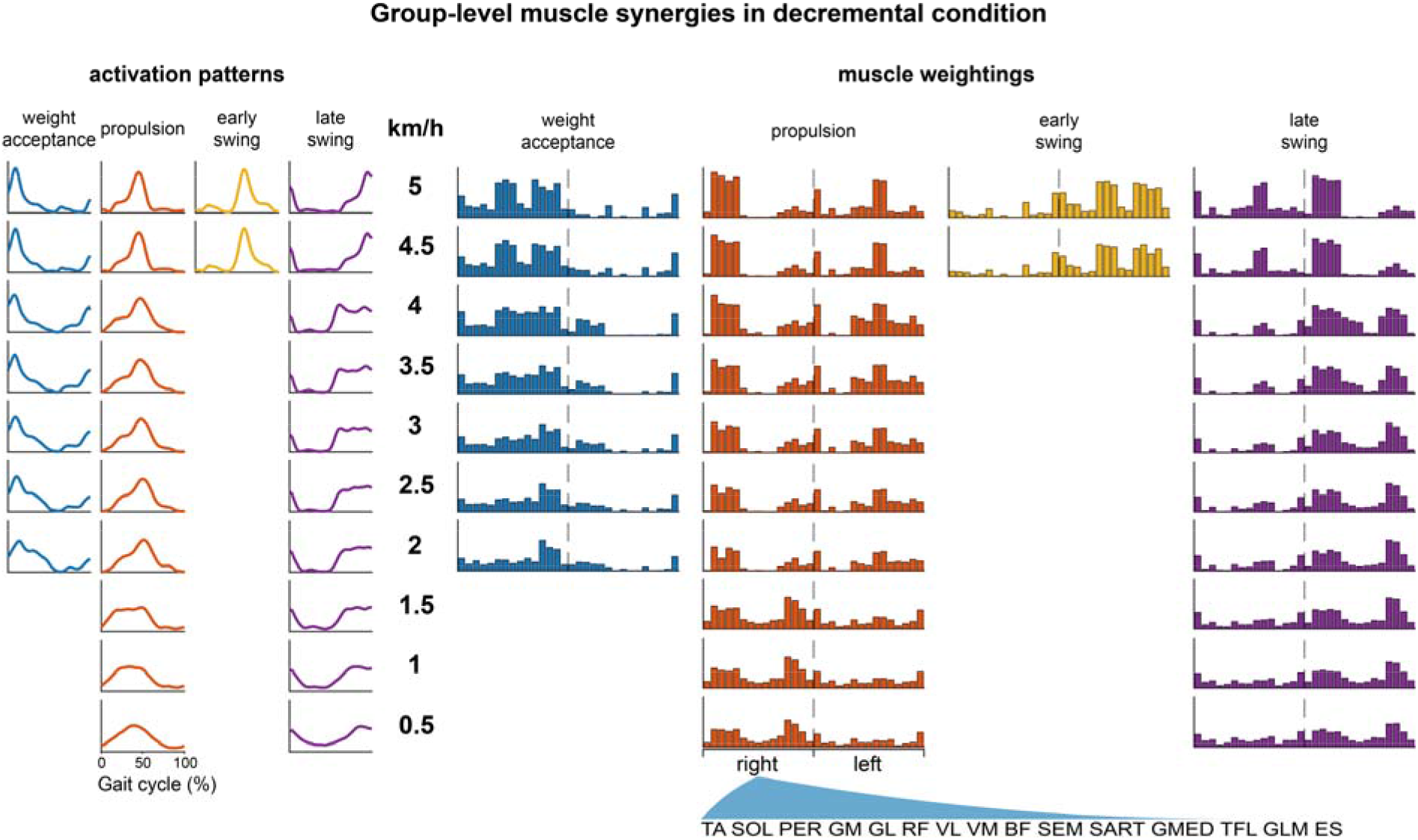
Group-level muscle synergies across speeds in the decremental condition. For each speed segment (5-0.5 km/h; rows), the figure shows the extracted synergy activation patterns (H; line plots on the left) and the corresponding muscle weightings (W; bar plot on the right). Activation patterns are time-normalized to the gait cycle (0-100%) and organized according to the functional phase of the gait cycle (weight acceptance, propulsion, early swing, and late swing). Bar plots show normalized muscle weights for the recorded muscles (labels in the bottom), with dashed vertical lines separating right- and left-side muscle groups. TA: tibialis anterior; SOL: soleus; PER: peroneus longus; GM: gastrocnemius medialis; GL: gastrocnemius lateralis; RF: rectus femoris; VL: vastus lateralis; VM: vastus medialis; BF: biceps femoris; SEM: semitendinosus; SART: sartorius; GMED: gluteus medius; TFL: tensor fasciae latae; GLM: gluteus maximus; ES: erector spinae.

In the incremental condition, low speeds (0.5–2 km/h) were adequately described by two synergies. From 2.5 km/h onward, an additional synergy emerged, yielding three synergies across intermediate speeds. At the highest speeds (4.5–5 km/h), a fourth synergy was identified (Figure 9).

In the decremental condition, four synergies were expressed at the highest speeds (5–4.5 km/h), after which the solution reduced to three synergies from 4 to 2 km/h. At the lowest speeds (≤1.5 km/h), the average modular organization further reduced to two synergies (Figure 10).

### Muscle synergy merging across walking speeds

Merging analysis revealed evidence of module merging across adjacent faster-to-slower speed-segment pairs, particularly at transitions where the number of synergies changed (Figure 11).

**Figure 11.**
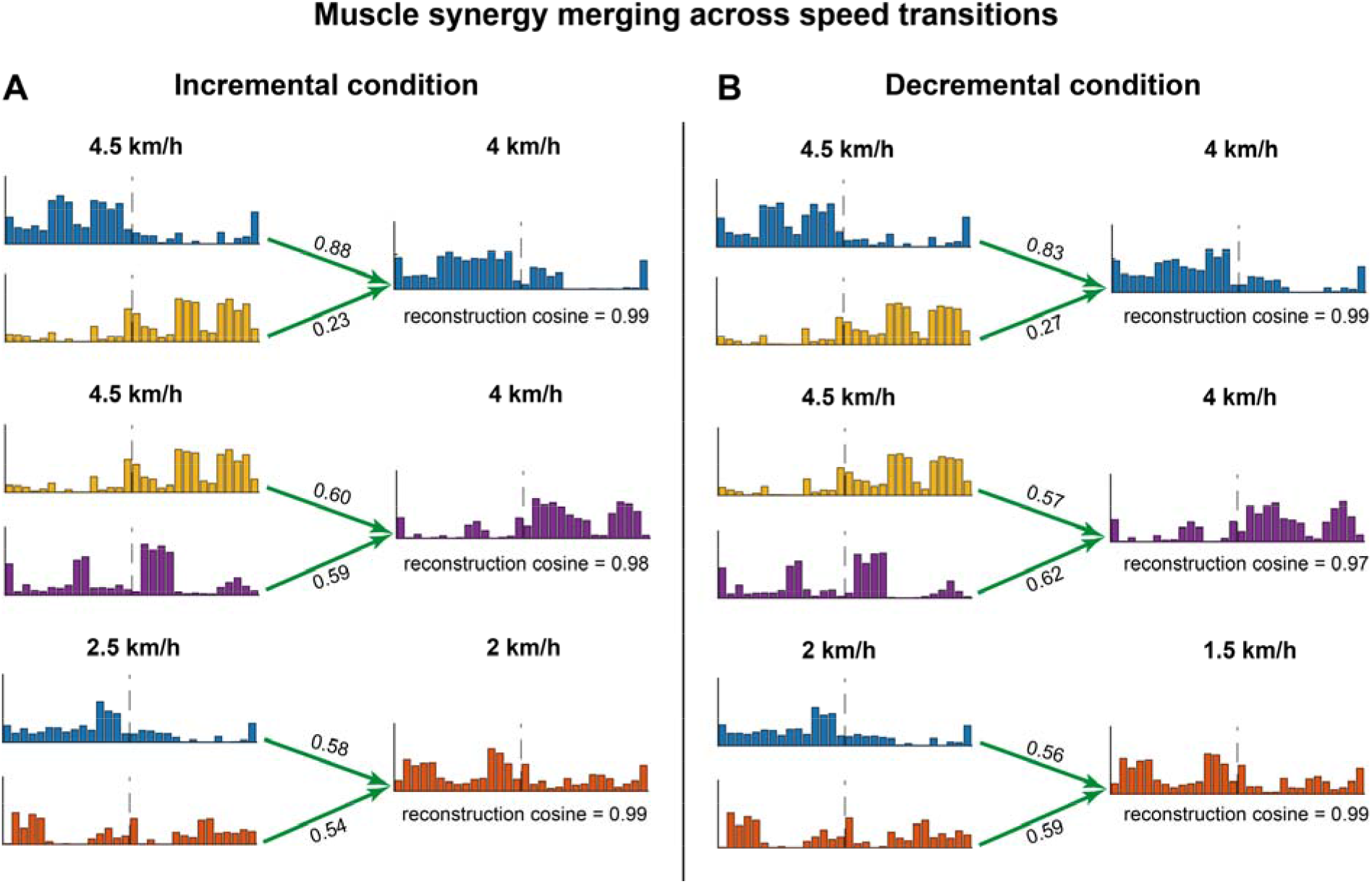
Examples of muscle synergy merging across speed transitions in incremental (A) and decremental (B) conditions. In each row, colored bar plots of muscle weightings (W) on the left show the contributing synergies from the adjacent faster speed segment, and the bar plot on the right shows the reconstructed target synergy in the slower segment. Green arrows indicate the contributing synergies used for reconstruction, and the values next to the arrows report the corresponding non-negative reconstruction coefficients (m_ik_). The overall reconstruction cosine similarity for each target synergy is reported below the right-hand panel. Examples illustrate merging at 2.5→2.0 km/h and 4.5→4.0 km/h in the incremental condition, and at 4.5→4.0 km/h and 2.0→1.5 km/h in the decremental condition.

In the incremental condition, merging was observed at 2 km/h ← 2.5 km/h for target synergy 1, reconstructed from synergies 1 (m_ik_ = 0.58) and 2 (m_ik_ = 0.54) (cosine = 0.99). A stronger merging pattern emerged at 4 km/h ← 4.5 km/h, where two target synergies were classified as merged: synergy 1 (cosine = 0.99), reconstructed from 1 and 3 (m_ik_ = 0.88, m_ik_ = 0.23) and synergy 3 (cosine = 0.98) from 3 and 4 (m_ik_ = 0.60, m_ik_ = 0.59) (Figure 11A).

In the decremental condition, merging was detected at 4 km/h ← 4.5 km/h for the same three targets (target 1 (cosine = 0.99) from synergies 1 and 3 (m_ik_ = 0.83, m_ik_ = 0.27), target 3 (cosine = 0.97) from synergies 3 and 4 (m_ik_ = 0.57, m_ik_ = 0.62), and at 1.5 km/h ← km/h for target 1 (cosine = 0.99) reconstructed from synergies 1 and 2 (m_ik_ = 0.56, m_ik_ = 0.59) (Figure 11B).

## Discussion

Overall, the present results show that walking speed modulates the balance between discrete- and rhythmic-like components of locomotor control. Movement smoothness metrics highlighted a transition from a more discrete-like organization at low speeds toward a more rhythmic regime at higher speeds. These findings suggest that the observed changes in smoothness cannot be explained by simple scaling of movement execution, but rather are consistent with a shift in control regime, from more segmented, discrete-like dynamics to more continuous, rhythmic organization. Specifically, across both incremental and decremental trials, LDJ and SPARC values became progressively less negative with increasing speed, indicating improved smoothness. Conversely, at the lowest speeds both metrics reached their most negative values, reflecting reduced smoothness.

These findings are consistent with literature suggesting that rhythmic behaviors are typically continuous and stereotyped, and thus smoother, whereas discrete actions introduce intermittency and higher-frequency fluctuations that increase jerkiness, thus reducing smoothness (Hogan and Sternad, 2007; Leconte et al., 2016; Park et al., 2017). Smoothness was quantified using LDJ and SPARC, which provide complementary time- and frequency-domain perspectives while mitigating limitations of conventional jerk estimates (Balasubramanian et al., 2012, 2015; Melendez-Calderon et al., 2021).

The speed-dependent smoothness pattern aligns with the dynamic-primitives framework, which proposes that very slow rhythmic movements are difficult to perform and can fragment into a sequence of discrete submovements (Park et al., 2017). Accordingly, extremely slow walking may reflect a more discrete-like control regime, whereas at higher speeds smoothness values stabilize, suggesting the emergence of a more stable rhythmic organization.

Notably, between-participant agreement decreased most markedly in an intermediate speed band (∼3.0-3.5 km/h), indicating a transition window in which participants showed heterogeneous responses before converging at faster walking speeds. These findings suggest that participants transition into a more stable rhythmic attractor at slightly different speeds, leading to reduced agreement within this intermediate range. At higher speeds, responses appeared to converge, suggesting stabilization within a predominantly rhythmic control regime (Flash and Hochner, 2005; Hogan and Sternad, 2013; Sternad et al., 2013). Furthermore, smoothness values showed higher between- and within-participant variability at the slowest speeds and stabilized with reduced variability at moderate-to-high speeds.

This is consistent with evidence that slow walking is associated with increased kinematic variability and systematic speed-dependent changes in local dynamic stability (e.g., Lyapunov-based measures) (Dingwell and Marin, 2006; England and Granata, 2007; McCrum et al., 2019). These findings support a nonlinear speed-smoothness relationship in which small increases at very slow walking speeds are associated with comparatively large changes in movement organization, whereas at moderate-to-fast speeds smoothness approaches stabilization and adjacent speed steps become less discriminable (Dingwell and Marin, 2006).

The present study also examined muscle synergy organization as a complementary measure of speed-dependent locomotor control. Subject-level muscle synergies exhibited a clear dependence on walking speed in both incremental and decremental conditions. At low speeds (0.5–1.5 km/h), activity was typically reconstructed with two synergies, increasing to three synergies at intermediate speeds and to four synergies at higher speeds. This aligns with evidence that human walking can be represented by a limited set of modules whose dimensionality varies with speed (Yokoyama et al., 2016). Similar modular structures have also been observed across development, with muscle synergies structuring locomotor patterns in children during both walking and running (Bach et al., 2021).

The incremental and decremental conditions suggested history dependence. During speed increases, dimensionality tended to rise stepwise (2 → 3 → 4), whereas during decreases the higher-dimensional solution persisted over a wider fast-speed range before reducing. Pairwise comparisons were consistent with this asymmetry: the first significant departure from the high-speed solution occurred at a higher speed during ramp-up than ramp-down, suggesting that slowing may initially be accommodated by retuning activations within an existing modular set before modules merge or drop out (Li, 2000; Hreljac et al., 2007).

Group-averaged results recapitulated the subject-level patterns. Also, synergy-merging analyses indicated that shifts in synergy number are concentrated at the transition speeds and are best explained by module merging. When stepping from a higher to an adjacent lower speed, the lower-speed synergies could be reconstructed with very high similarity as linear combinations of the higher-speed synergies, implying that the reduction in dimensionality reflects merging rather than module elimination. Our results are consistent with synergy-merging literature, where altered or lower-dimensional synergy sets can be explained as combinations of a higher-dimensional reference set (Cheung et al., 2020, 2012; Hashiguchi et al., 2016; Mizuta et al., 2022).

Overall, kinematic and EMG results showed coherent evidence of speed-dependent reorganization characterized by a low-speed regime with reduced smoothness and increased variability, an intermediate transition window with higher heterogeneity across participants, and a high-speed regime in which both smoothness and synergy number stabilized. speed This profile supports the interpretation that walking speed modulates the balance between discrete- and rhythmic-like control. Increasing speed was associated with smoother, less variable kinematics and a richer modular organization of muscle activity, consistent with a more stable rhythmic organization. Conversely, as speed decreased, smoothness deteriorated and inter-individual variability increased, while EMG organization showed a reduction in the number of synergies, consistent with a merging of modules, suggesting a shift toward a more discrete-like regime.

A few limitations should be considered. First, some EMG channels were excluded in certain participants due to signal-quality criteria (e.g., noise, motion artifacts, electrode issues), which may influence absolute dimensionality estimates, although the main speed-dependent patterns remained evident. Second, treadmill walking differs from overground gait in sensory context and spatiotemporal characteristics; therefore, caution is warranted when generalizing absolute values. Nonetheless, the treadmill enabled systematic speed manipulation under controlled conditions. Future work should replicate these analyses overground and with broader EMG coverage where feasible.

More broadly, these findings support the utility of the dynamic-primitives framework for interpreting speed-dependent locomotor organization, with potential implications for motor learning and rehabilitation. This framework may inform therapeutic strategies (Leconte et al., 2016) and improve our understanding of motor learning (Hogan et al., 2006). Hogan and Sternad, (2013) proposed that motor learning and recovery involve encoding dynamic primitive parameters to reconstruct movement patterns rather than reproducing specific behaviors, as seen in the progression of infant reaching from discrete submovements to continuous, rhythmic patterns (Berthier, 1996; Von Hofsten, 1991). Similar patterns have been observed in stroke recovery, where early stereotyped submovements gradually become larger, fewer, and more integrated (Krebs et al., 1999; Rohrer et al., 2002; Dipietro et al., 2009) although it remains unclear whether similar mechanisms apply to lower limb locomotion.

## Acknowledgments

This work was supported by European Research Council (ERC) under the European Union’s Horizon 2020 research and innovation program (grant agreement n: 715945 Learn2Walk), and by the Dutch Organization for Scientific Research (NWO) VIDI grant (grant agreement n: 016.156.346 FirSTeps), which supported the acquisition of the experimental instrumentation. We would like to thank Pascal Thom Sprenger for his support during data collection.

## Author contributions

Conceptualization: G.P., R.B., D.M., N.D.; Methodology: G.P., N.D.; Formal analysis: G.P.; Investigation: G.P, N.D.; Data Curation: G.P.; Writing – original draft: G.P.; Writing Review & Editing: R.B, D.M., N.D.; Visualization: G.P.; Supervision: R.B., D.M., N.D.

