## Supplementary Table 1 for "Speed-driven transitions between discrete and rhythmic dynamics in walking revealed by kinematic smoothness and muscle synergies"

| **Subjects** | **Age**  **(years)** | **Gender** | **Weight**  **(kg)** | **Height**  **(cm)** |
| --- | --- | --- | --- | --- |
| P1 | 21 | F | 64 | 168 |
| P2 | 21 | F | 60 | 166 |
| P3 | 23 | M | 80 | 190 |
| P4 | 22 | M | 80 | 180 |
| P5 | 27 | F | 48 | 160 |
| P6 | 22 | M | 86 | 186 |
| P7 | 22 | M | 72 | 179 |
| P8 | 22 | M | 80 | 183 |
| P9 | 18 | F | 72 | 182 |
| P10 | 25 | M | 79 | 175 |
| P11 | 22 | F | 55 | 167 |
| P12 | 20 | F | 53 | 163 |
| P13 | 19 | F | 54 | 166 |
| P14 | 17 | F | 53 | 162 |
| P15 | 19 | F | 66 | 171 |
| P16 | 20 | F | 56 | 169 |
| P17 | 19 | F | 59 | 165 |
| P18 | 19 | F | 64 | 172 |

**Table S1. Participant Characteristics**
